# Morphological control of receptor-mediated endocytosis

**DOI:** 10.1101/2021.10.01.462837

**Authors:** Daniele Agostinelli, Gwynn J. Elfring, Mattia Bacca

**Author notes:** Corresponding author. Email address (Mattia Bacca).

## Abstract

Receptor-mediated endocytosis is the primary process for nanoparticle uptake in cells and one of the main entry mechanisms for viral infection. The cell membrane adheres to the particle (nanoparticle or virus) and then wraps it to form a vesicle delivered to the cytosol. Previous findings identified a minimum radius for a spherical particle below which endocytosis cannot occur. This is due to the insufficient driving force, from receptor-ligand affinity, to overcome the energy barrier created by membrane bending. In this paper, we extend this result to the case of clathrin-mediated endocytosis, which is the most common pathway for virus entry. Moreover, we investigate the effect of ligand inhibitors on the particle surface, motivated by viral antibodies, peptides or phage capsids nanoparticles. We determine the necessary conditions for endocytosis by considering the additional energy barrier due to the membrane bending to wrap such inhibiting protrusions. We find that the density and size of inhibitors determine the size range of internalized particles, and endocytosis is completely blocked above critical thresholds. The assembly of a clathrin coat with a spontaneous curvature increases the energy barrier and sets a maximum particle size (in agreement with experimental observations on smooth particles). Our investigation suggests that morphological considerations can inform the optimal design of neutralizing viral antibodies and new strategies for targeted nanomedicine.

## Introduction

Endocytosis is a fundamental process by which cells uptake external particles. This can occur through different pathways (Yi et al., 2014). One observed mechanism exploits the binding affinity between cell receptors and particle ligands, leading the cell membrane to wrap the whole particle into a delivery vesicle. Another mechanism involves proteins - present in the cytoplasm - which form a coated vesicle having spontaneous curvature, thereby facilitating the internalization of particles with selected size (Mashl and Bruinsma, 1998).

Cells commonly use endocytosis to internalize nutrients, but the process can also result in undesired viral entry and hence the infection of the cell. A physical description of endocytosis is thus crucial for understanding nutrient uptake, viral infection, and nanoparticle engineering for diagnostics and targeted therapies requiring a selective uptake.

Receptor-mediated endocytosis depends on particle size and geometry, biochemistry, and mechanics (Zhang et al., 2015). Previous studies considered simple particle shapes such as spheres and cylinders, and showed that endocytosis requires the particle to be larger than a minimum size (Gao et al., 2005; Yi et al., 2014; Yi and Gao, 2017). However, these results neglected the role of coating proteins such as clathrin and caveolin, despite the fact that clahtrin-mediated endocytosis is the most common entry process of viruses (Marsh and Helenius, 2006; Flint et al., 2015). Moreover, the effect of inhibiting protrusions on endocytosis remains largely unexplored. In particular, it is unknown how ligand inhibitors quantitatively affect particle uptake by their twofold action of reducing the ligand density and modifying the particle morphology.

We consider a spherical and rigid particle having a fraction of ligands blocked by finite-sized inhibitors that protrude out of the spherical surface. The inhibitors represent neutralizing antibodies, peptides, or phage capsids attaching to the surface of the particle to cover (neutralize) a fraction of ligands (Flint et al., 2015; Lauster et al., 2020). The physical scale of the inhibitors adds an additional geometric consideration for the successful encapsulation of the particle. We focus on the spontaneity of the wrapping process initiating endocytosis and explore the geometric determinants of the complex morphology obtained. The number and size of protrusions raise the energetic barrier for endocytosis based on particle size and inhibitor spacing due to both the reduction of ligand-receptor bindings and the additional bending penalty to wrap the inhibitors. Fig. 1 illustrates the concept of inhibiting protrusions that block the particle entry.

**Figure 1:**
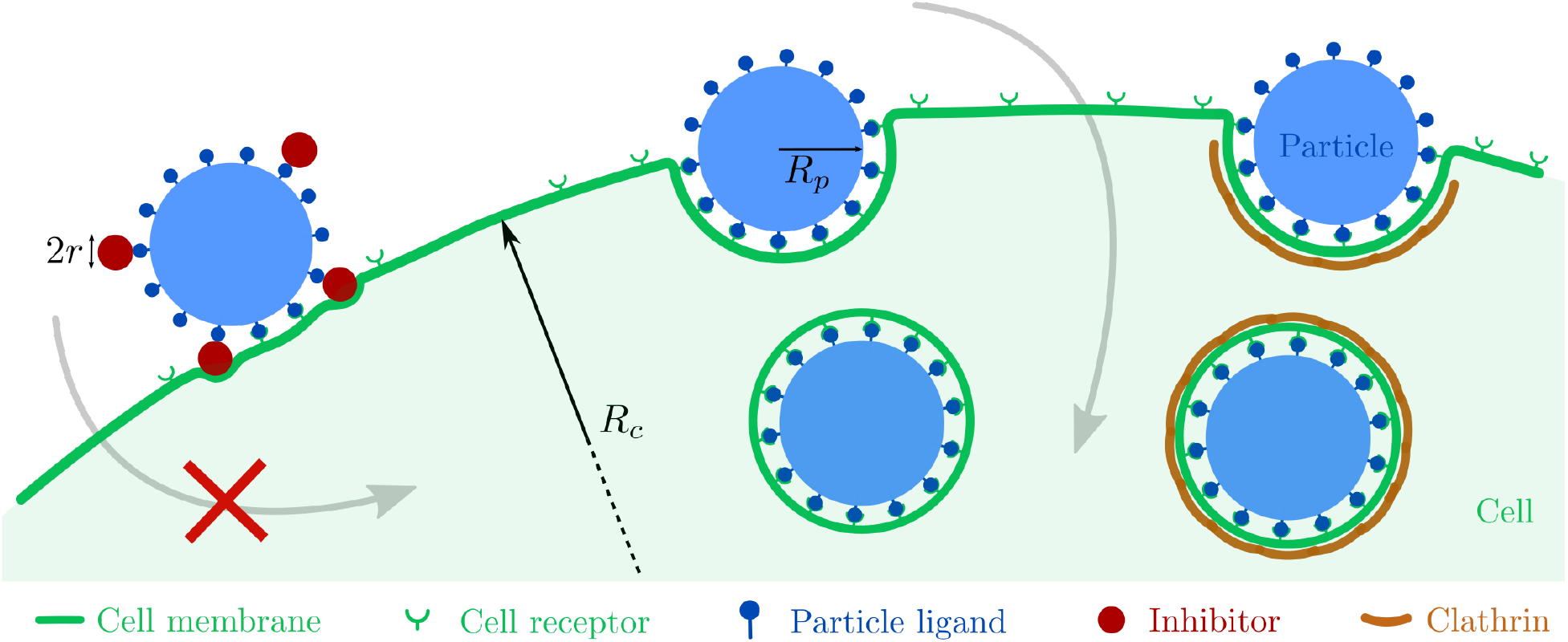
Schematic illustration of the process of receptor-mediated endocytosis, including the case of clathrin coating. The presence of ligand inhibitors on the surface of a spherical particle can prevent cell entry, depending on the size and number of the inhibitors. To reduce the number of variables, we assume the cell radius *R*_*c*_ to be much larger than that of the particle *R*_*p*_ (infinite and flat cell membrane).

Building on the mechanical model for the uptake of a rigid sphere (Gao et al., 2005), we account for the diffusivity of cell receptors across the cell membrane and the free energy of the membrane as the sum of elastic deformation energy and adhesion energy. The spontaneity of endocytosis is determined by imposing that the rate of free energy reduction, due to membrane wrapping, equals the rate of free energy dissipation due to receptor transport. We account for the reduced ligand density due to the presence of inhibitors and calculate the equilibrium configuration of the membrane wrapping around the particle and its protrusions.

We find that size, indentation, and density of ligand inhibitors determine the selective uptake of particles, and the entry is completely blocked above critical thresholds. In particular, when endocytosis is independent of protein coats, particles can enter the cell only if their radius is larger than a minimum value, while for the clathrin-mediated endocytosis, particles are selected within a size range, consistently with experiments on spherical particles (Rejman et al., 2004).

The proposed results provide a foundation for the design of viral antibodies, peptides, and capsids acting as physical and geometrical inhibitors, and for the design of targeted nanoparticles undergoing selective cell entry based on the interplay between morphology and receptor density.

## Theory

### Particle geometry

We consider a spherical particle of radius *R*_*p*_ with ligands uniformly distributed on its surface with a density *ξ*_*L*_. We estimate the spacing between two adjacent ligands 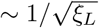 by assuming an ideal tessellation of the sphere consisting of a large number of hexagons and 12 pentagons (such as in Goldberg polyhedrons), see Appendix A. We denote by *R* the curvature radius of the membrane that wraps the spherical particle, thus including the particle radius *R*_*p*_, the receptor-ligand bond, and half thickness of the lipid bilayer (1.5− 2.5 nm, Lewis and Engelman (1983)), see Fig. 2a. In the following we refer to *R* as the particle radius. The wrapping process requires cell receptors to diffuse towards the contact region so that the receptor density on the cell surface approximates the ligand density on the particle surface, *ξ*_*L*_.

**Figure 2:**
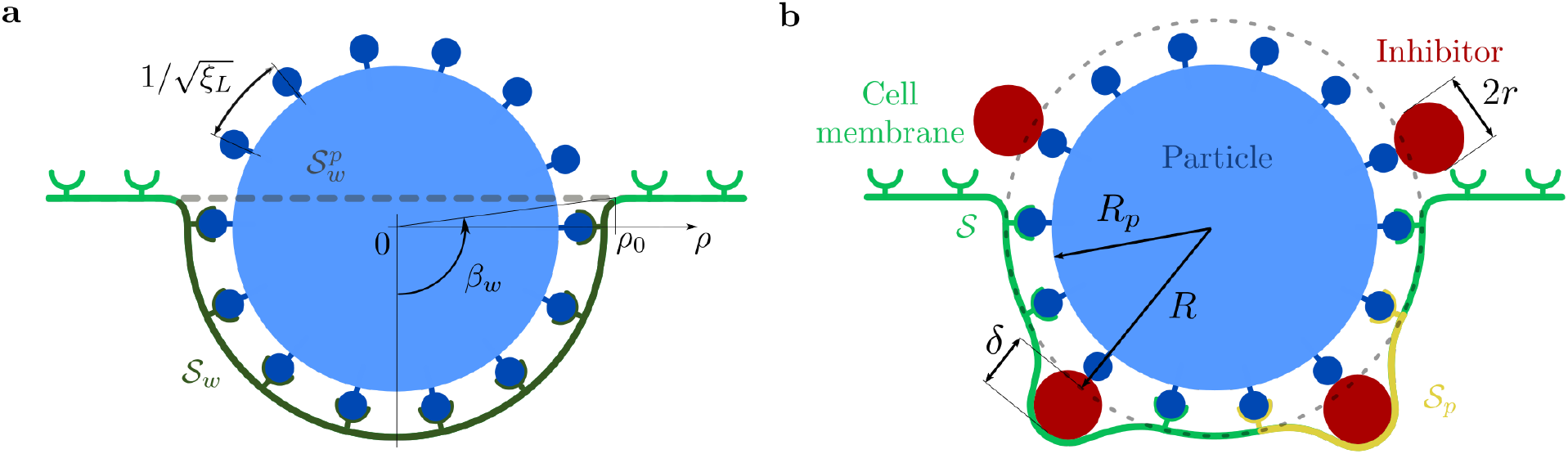
Two-dimensional schematic illustration of the cell membrane 𝒮 that wraps an external particle during receptormediated endocytosis. **(a)** Spherical particle of radius *R*_*p*_ with ligand surface density *ξ*_*L*_, so that the spacing between two adjacent ligands is approximately 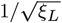, see Appendix A. The cell membrane develops a curvature of radius *R*, which includes both the particle and the receptor-ligand bonds. 𝒮_*w*_ denotes the portion of the cell membrane in contact with the particle, which corresponds to a wrapping angle *β*_*w*_. 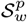 is the projection of 𝒮_*w*_ onto the flat membrane. **(b)** A fraction *p* of ligands are inhibited by spherical protrusions of radius *r*. Inhibitors protrude a length *δ*, and 𝒮_*p*_ denotes the representative portion of cell membrane that wraps a single protrusion, assuming protrusions are largely spaced.

We cover a fraction *p* of ligands with spherical inhibitors of radius *r*, which we assume uniformly distributed on the particle. Such inhibitors protrude over the particle surface a length *δ* (see Fig. 2b), which we express as a multiple of the inhibitor radius *r, i*.*e*., *δ* := 2*dr* where *d* is a dimensionless indentation parameter.

When the cell membrane wraps the particle, including its protrusions, the receptors diffuse toward the wrapping region to bind to the particle ligands that are still active. The surface density of effective ligands is reduced to (1 − *p*)*ξ*_*L*_, while the membrane will need additional area and bending to wrap around the protrusions. The surface density of receptors in the wrapped membrane is

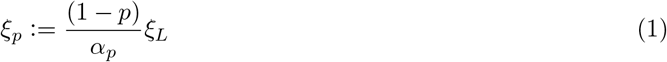

where *α*_*p*_ := *A*_tot_*/A*_*s*_ is the dimensionless ratio between the surface area of the membrane wrapping the whole particle with inhibiting protrusions, *A*_tot_, and the surface area of the spherical particle, *A*_*s*_ = 4*πR*^2^. *α*_*p*_ ≥ 1 measures the excess area needed for wrapping that is due to the inhibitors, with *α*_*p*_ = 1 when *p* = 0 (no protrusions).

The parameter *p* defines the fraction of inhibited ligands and we assume that there is a maximum value *p*_max_ ≈ 1*/π*, for which the wrapping process is still possible in a symmetric fashion. Indeed, above this threshold value, the number of inhibitors is so large that there must be at least two adjacent inhibitors, so that we cannot assume that the membrane wraps any protrusion by binding to the surrounding ligands, see Appendix B.

### Free energy of the cell membrane and protein coat

We consider a membrane 𝒮 with an initially uniform receptor density *ξ*_0_, which curves to wrap a proximal particle, as cell receptors bind to particle ligands. We assume the membrane to be infinitely extended and initially flat, *i*.*e*. the particle radius *R*_*p*_ is much smaller than the cell radius *R*_*c*_, namely, *R*_*p*_*/R*_*c*_ ≪ 1 (see Fig. 1).

Since endocytosis is often mediated by proteins - such as clathrin and caveolin - that form coated pits (Flint et al., 2015), we include a protein coat with a spontaneous mean curvature *H*^⋆^, which assembles during the wrapping process and attaches to the membrane in the contact region.

The free energy of the cell membrane is given by the contributions from elastic deformation *ε*_el_ and adhesion *ε*_ad_, namely, *ε* = *ε*_el_ + *ε*_ad_. We describe *ε*_el_ with the Canham-Helfrich model (Canham, 1970; Helfrich, 1973) and neglect deformations of the cell membrane outside the contact region 𝒮_*w*_, such that

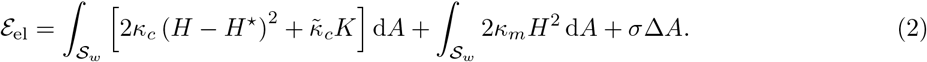

In Eq. (2), the first integral term provides the elastic energy of the protein coat (denoted by the subscript c), whereas the other two terms provide the elastic energy of the cell membrane (denoted by the subscript m). The bending energy is expressed in terms of the mean and Gaussian curvatures, respectively *H* and *K*, with the corresponding elastic moduli *κ* and 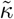. The Gaussian curvature accounts for the energy stored in symmetric saddles with zero mean curvature. In view of the Gauss-Bonnet theorem, its integral over the surface reduces to the sum of a topological invariant and the integral of the geodesic curvature of the surface boundary; we neglect this term (a constant) for the infinite membrane and we consider it only for the protein coat, whose boundary evolves during wrapping. For simplicity, we assume that the spontaneous mean curvature of the protein coat, *H*^⋆^, is constant even though recent experimental and theoretical evidence supports a flat-to-curved transition during the assembly of clathrin pits (Frey, 2019). Moreover, we consider that both the membrane and the coat have the same configuration (*i*.*e*., curvatures). This hypothesis implies that we neglect the coat thickness, which is about 5 nm for caveolin (Stoeber et al., 2016), and 20 − 30 nm for clathrin (Vigers et al., 1986; Pedersen, 1993). The last term in Eq. (2) describes the mechanical work due to membrane stretching, *i*.*e*. the product of surface tension *σ* to the excess area 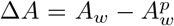. *A*_*w*_ is the area of the wrapping surface 𝒮_*w*_, and 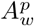 is the area of the corresponding projection 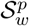 on the flat configuration (Deserno, 2004), see Fig. 2.

The adhesion energy *ε*_ad_ accounts for entropic, *ε*_r_, and enthalpic, *ε*_h_, contributions (Zhang et al., 2015). The configurational, entropic, energy of receptors (in analogy with an ideal gas, Freund and Lin (2004)) is

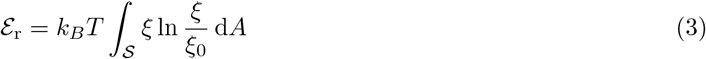

where *k*_*B*_ is the Boltzmann constant, *T* is the absolute temperature, *ξ* is the number of receptors per unit area of the cell membrane, and *ξ*_0_ is the uniform receptor density in an isolated cell, or at infinite distance from the wrapping site.

The change of enthalpy, due to the formation of coated pits and the receptor-ligand binding (Gao et al., 2005), provides the driving force for the wrapping process and is given by

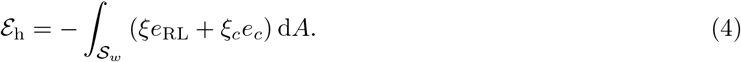

Here *e*_RL_ is the receptor-ligand binding energy, *ξ*_*c*_ is the surface density of coating protein bonds, and *e*_*c*_ is the protein binding energy.

Let us now divide each energetic contribution by *k*_*B*_*T*, and each length by the ligand spacing 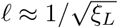. The dimensionless free energy is then

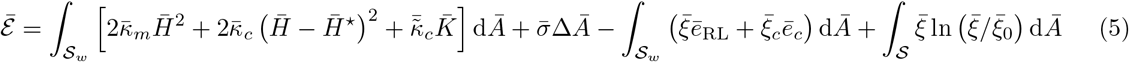

where 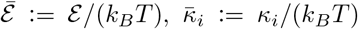 for 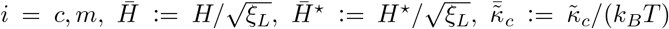, 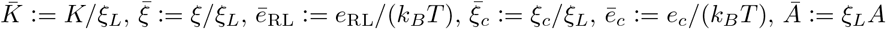, and 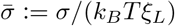. In Table 1 we report the physical quantities adopted in our model, taken from literature.

**Table 1:** Relevant values of the model parameters as reported in the biological literature. The thermal energy constant is *k*_*B*_*T* ≈ 4.14 pN nm (at a temperature of *T* = 300K).

| Parameter | Description | Value/Range | Units | Reference(s) | Used value(s) |
| --- | --- | --- | --- | --- | --- |
| $\kappa_m$ | Membrane bending modulus | 10 – 25 | $k_B T$ | Steigmann (2017), Banerjee et al. (2012), Gao et al. (2005) | $\bar{\kappa}_m = 20$ |
| $\kappa_c$ | Clathrin coat bending modulus | 300 | $k_B T$ | Saleem et al. (2015), Nossal (2001) | $\bar{\kappa}_c = 300$ |
| $\tilde{\kappa}_c$ | Saddle-splay modulus of clathrin coat | n.a. | $k_B T$ | n.a. | $\tilde{\kappa}_c = 0$ |
| $\xi_0$ | Cell receptor density | $5 \cdot 10^{-5} - 130 \cdot 10^{-5}$ | $\text{nm}^{-2}$ | Gao et al. (2005), Yi and Gao (2017) | $\bar{\xi}_0 = 0.025, 0.1$ |
| $\xi_L$ | Particle ligand density | $3 \cdot 10^{-3} - 20 \cdot 10^{-3}$ | $\text{nm}^{-2}$ | Gao et al. (2005), Yi and Gao (2017) | $\xi_L = 5 \cdot 10^{-3}$ |
| $e_{RL}$ | Receptor-ligand binding energy | 10 – 25 | $k_B T$ | Yi and Gao (2017) | $\bar{e}_{RL} = 15$ |
| $e_c$ | Binding energy per clathrin triskelion | 5 – 30 | $k_B T$ | Saleem et al. (2015), Banerjee et al. (2012), Kumar and Sain (2016) | $\bar{e}_c = 23$ |
| $\xi_c$ | Density of clathrin triskelia | $1.25 \cdot 10^{-3}$ | $\text{nm}^{-2}$ | Saleem et al. (2015) | $\bar{\xi}_c = 0.25$ |
| $\sigma$ | Membrane tension | $7 \cdot 10^{-4} - 2.93$ | $\frac{k_B T}{\text{nm}^2}$ | Morris and Homann (2001), Derényi et al. (2002), Deserno (2004), Frey (2019) | $\bar{\sigma} = 1$ |
| $R^*$ | Preferred radius of clathrin-coated pit | 32.5 – 50 | nm | Heuser (1980), Traub (2011), Saleem et al. (2015) | $\bar{H}^* = -0.283$ |

In clathrin-mediated endocytosis, the self-assembling coat controls the equilibrium configuration of the bilayer - composed of membrane and coat - wrapped around the protrusion. This is due to the higher bending rigidity of the coat 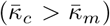, as shown in Table 1. If either the coat is absent or the membrane accommodates to the spontaneous curvature of the coat, then the elastic term is dominated by the cost of bending the cell membrane, since its rigidity, 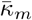, is one order of magnitude larger than its tension, 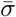, see Table 1. The adhesion energy is of the same order of magnitude of the elastic energy, with the enthalpic term dominating over the entropic one. Indeed, in the wrapping region, 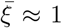 is one order of magnitude smaller than the binding energies *ē*_*RL*_ and *ē*_*c*_. From this analysis we observe that the process is primarily led by an exchange of enthalpic (binding) energy, 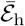, with elastic (wrapping) energy, 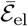.

In comparison with a spherical particle, the presence of protrusions hinders the process of endocytosis by increasing the energetic cost of wrapping. Each protrusion raises *ε*_el_ by increasing membrane curvature locally. Protrusions also affect the adhesion energy *ε*_ad_ by reducing the enthalpic gain and the entropic cost, due to the loss of a fraction *p* of receptors in the wrapped region. This can be seen from *ξ*_*p*_ < *ξ*_*L*_ in Eq. (1) and the last two terms of Eq. (5). The reduced cost favors endocytosis, but is lower than the reduced gain, which finally provides an additional barrier to particle uptake. In the following section we provide quantitative evidence of this scaling argument.

### Condition for endocytosis

#### Spherical particle

We assume that cell receptors are mobile and diffuse towards the contact site to bind to (immobile) particle ligands. We assume that receptors are uniformly distributed with a surface density *ξ*_0_ at remote distances from the binding site. They accumulate in the contact region to match the surface density of ligands, *ξ*_*L*_ > *ξ*_0_. This yields a depletion of receptors in the outer proximity of the contact front, which determines a concentration gradient that drives the global diffusion towards the binding site (Freund and Lin, 2004; Gao et al., 2005).

We describe the nonuniform receptor density outside the contact region with the function *ξ*(*ρ, t*), where *ρ* is the radial coordinate and *t* is time. We denote by *ρ*_0_(*t*) the radial coordinate of the moving contact front, which is given by *ρ*_0_(*t*) = *R* sin *β*_*w*_(*t*), where *β*_*w*_(*t*) is the angle that corresponds to the surface area *A*_*w*_(*t*) of the portion of the cell membrane that wraps the particle, 𝒮_*w*_(*t*) (see Fig. 2a).

During the wrapping process, receptors are driven by a local reduction in the free energy, which rate of change is

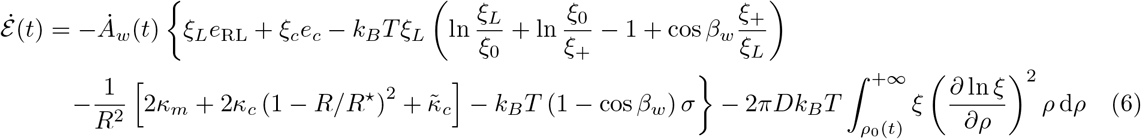

as explained in Appendix C with *p* = 0. Here a superposed dot denotes time differentiation, *D* is the diffusivity of cell receptors, *R*^⋆^ is the preferred radius of curvature of the protein coat, and 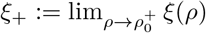 is the receptor density in the unwrapped region next to the binding site 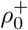, giving *ξ*_+_ < *ξ*_0_ (Freund and Lin, 2004).

The integral term in Eq. (6) is the rate of energy dissipation due to receptor transport (Freund and Lin, 2004). The expression within {…} describes the rate of free energy change per unit of new wrapping membrane. The first two terms represent the reduction of enthalpy (receptor-ligand bindings and coat assembling). The expression within (…), and multiplied by *ξ*_*L*_, accounts for the entropic change: the first two terms describe the relocation of receptors from the uniform density *ξ*_0_ to *ξ*_*L*_, within the wrapped region, and *ξ*_+_, outside that region; the other two terms describe the receptor transport across the front, from *ξ*_+_ to *ξ*_*L*_, where *ξ*_+_ is multiplied by cos *β*_*w*_ to account for the difference between the spherical area and its planar projection. The terms within […] define the free energy increment required to deform elastically the membrane and the protein coat. Finally, the term multiplied by *σ* accounts for the membrane tension, which increases with the wrapping angle *β*_*w*_.

Particle wrapping can occur only if the free energy is released at a rate greater than or equal to the rate of energy dissipated by the cell receptor transport across the membrane (Freund and Lin, 2004). From this, we determine the necessary condition for initiating particle wrapping via Eq. (6), by approximating the density *ξ*_+_ with the initial density *ξ*_0_, and by setting *β*_*w*_ = 0, giving

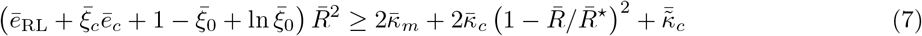

with 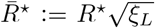. If we neglect the protein coat, Eq. (7) reduces to the condition found by Gao et al. (2005). In this case, if the expression within the parenthesis on the left side is positive - *i*.*e*., the enthalpic gain is higher than the entropic cost - the inequality is satisfied for any value of the dimensionless radius larger than a threshold, 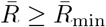. By fixing the value of the ligand density, this translates to the existence of a minimum radius *R*_min_ for the particle uptake. For protein-independent receptor-mediated endocytosis and for relevant biological values of the parameters (see Table 1), the minimum radius is *R*_min_ ≈ 24 nm, which is in agreement with experimental observations (Gao et al., 2005).

In the presence of a protein coat, the same holds true for values of the preferred radius of curvature that are higher than a threshold 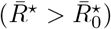, such that for particles that are large enough, the driving force is sufficient to overcome the penalty for deviating from 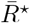. However, for higher values of the spontaneous curvature of the protein coat 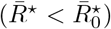, the cost of deviating from the preferred configuration is increasing faster than the driving force, for particles of increasing radii. This implies also an upper bound on the dimensionless radius, 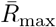. In other terms, if the protein coat has a preferred radius smaller than 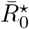, the cost of bending dominates over adhesion and becomes prohibitive for radii larger than a maximum value. In this case, the wrapping is favorable only for particles within a specific size range, 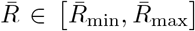, see Appendix D.

For relevant values of the biological parameters for clathrin-mediated endocytosis (see Table 1), the critical preferred radius is 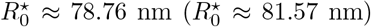 for 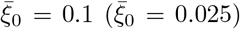, cf. Appendix D. Thus, by assuming a preferred radius of curvature *R*^⋆^ = 50 nm, we predict the uptake of particles of radius between *R*_min_ ≈ 33 nm (*R*_min_ ≈ 34 nm) and *R*_max_ ≈ 134 nm (*R*_max_ ≈ 126 nm) for 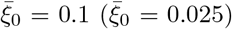. This theoretical conclusion is in good agreement with experimental observations about the size dependence of clathrin-mediated endocytosis that exhibited an upper radius limit for internalization of approximately 100 nm (Rejman et al., 2004).

#### Spherical particle with protruding inhibitors

We adapt the argument sketched above for the spherical particle, by splitting the cell membrane into regions that wrap a single protrusion and the rest that adheres to the spherical part of the particle, see Fig. 2b. As shown in Appendix C, we arrive at the inequality

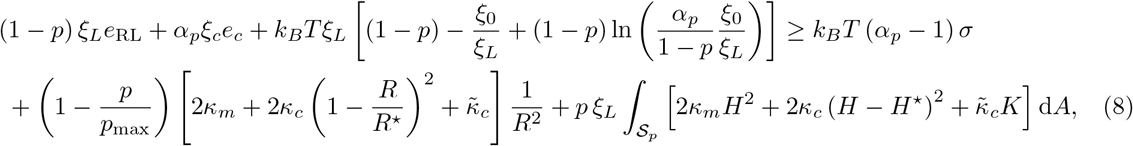

where 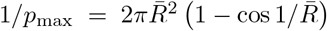, see Appendix B. Eq. (8) provides a necessary condition for the initiation of wrapping for a particle with inhibiting protrusions, and reduces to Eq. (7) for *p* = 0.

To solve Eq. (8) we need to obtain the equilibrium configuration of the membrane in the region that wraps a protruding inhibitor, 𝒮_*p*_. This reduces to the solution of an obstacle problem that implicitly depends on the geometry of the particle, namely, *r* (protrusion radius), *R*, and *δ* (indentation), see Fig. 2b. The equilibrium configuration is the one that minimizes the elastic deformation energy, *ε*_el_, while satisfying compatibility with the protrusion and adhered to the particle at the boundary, due to the surrounding receptor-ligand bonds. This requires the solution of a nonlinear variational inequality, which we approximate numerically, since analytical results are unavailable. In this study we used the finite element method to minimize the elastic energy with a penalty term that enforces the obstacle constraint, described as a half spherocylinder in the Monge parametrization, see Appendix E and Fig. 3.

**Figure 3:**
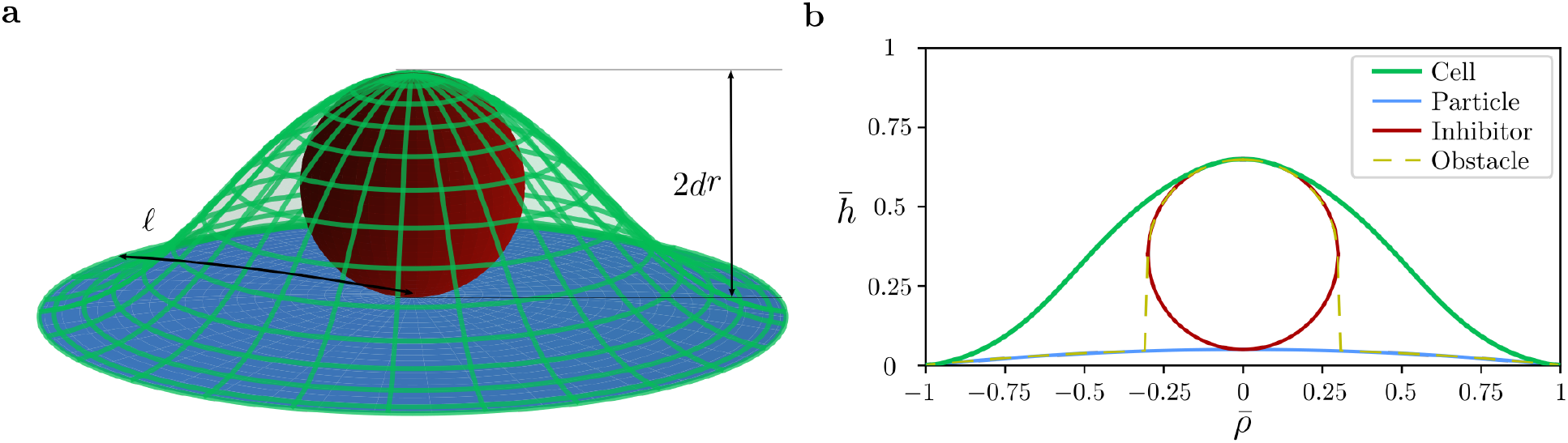
**(a)** Three-dimensional and **(b)** two-dimensional views of cell membrane clamped to the particle at the boundary and wrapping the ligand inhibitor. Both the radial coordinate 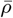, and the height function 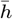 are measured in units of the ligand spacing 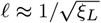. We determined the configuration as an axisymmetric solution to the obstacle problem where the obstacle is defined as half capsule (yellow dashed line), for model parameters *ℓ* ≈ 14 nm, 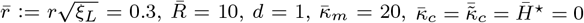, see Appendix E.

In this analysis, we assumed that the membrane wraps the protrusions and binds to the closest effective ligands. This hypothesis is realistic for small *p* and small *r*, where a single-spaced configuration is energetically favorable. In fact, for larger *r*, a doubled-spaced configuration is favored over the single-spaced one (see Appendix F).

## Results

By solving for the equality in Eq. (8) in the parameter space 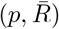 we determine the regions where endocytosis is favorable, using the parameters reported in Table 1. Fig. 4a shows the results in the absence of clathrin coat, whereas, Fig. 4b shows the results in the presence of the coat.

**Figure 4:**
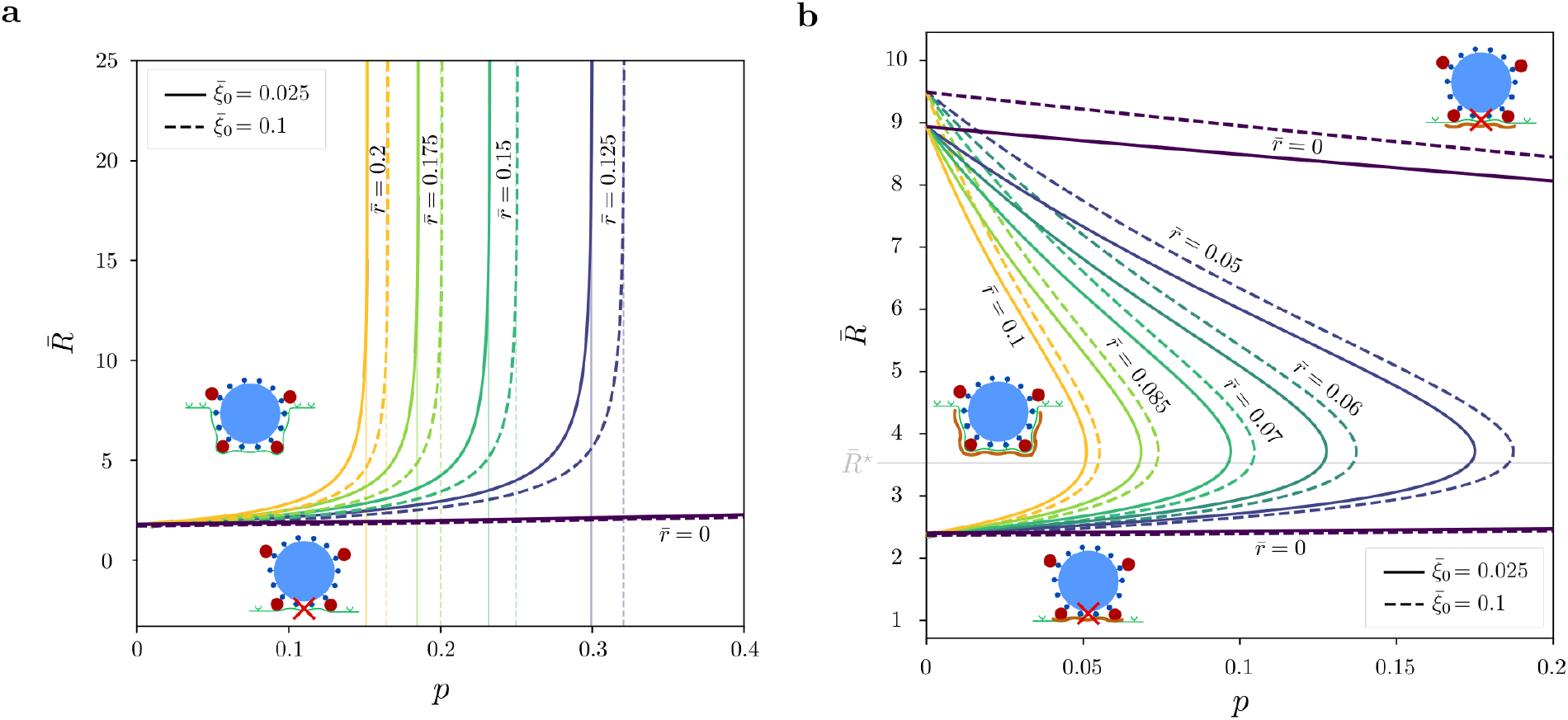
Regions of favorable endocytosis in the parameter space 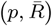 where *p* is the fraction of inhibited ligands and 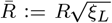 is the dimensionless particle radius. Region boundaries are determined from Eq. (8) for different dimensionless obstacle sizes, 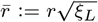, and ratios between ligand and receptor densities, 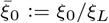. **(a)** Receptor-mediated endocytosis without protein coat for model parameters *d* = 1, *ē*_RL_ = 15, 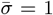, *ξ*_*L*_ = 5 ·10^−3^ nm^−2^, and 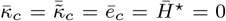. The curve provides the minimum radius 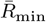 and endocytosis is favored when 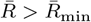. Vertical lines show the critical values of *p* that correspond to the limit case of 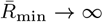. **(b)** Clathrin-mediated endocytosis for model parameters *d* = 1, *ē*_RL_ = 15, 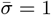, *ξ*_*L*_ = 5 ·10^−3^ nm^−2^, 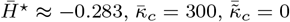, and *ē*_*c*_ = 23. The curve defines the minimum and maximum radii, 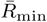 and 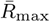, and endocytosis is favored when 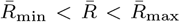. The horizontal (grey) line reports the dimensionless radius of spontaneous curvature of the clathrin coat 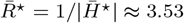.

In the absence of inhibitors (*p* = 0), the wrapping process starts only if the particle radius lies within the specific range *R*_min_ ≤ *R* ≤ *R*_max_, with *R*_max_ → ∞ in the absence of protein coat. In the case of inhibitors (*p >* 0) without protrusions 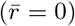, the range size [*R*_min_, *R*_max_] slightly contracts for moderate values of *p* (see Fig. 4). Finally, the presence of protruding inhibitors (*p >* 0 and 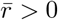) significantly shrinks the range of admissible radii until their density *p* reaches a critical value 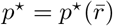, above which the particle uptake is completely blocked. This shows the crucial impact of the inhibitor size 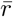 for moderate values of the fraction of inhibited ligands *p*. As observed in Fig. 4, *p*^⋆^ is inversely related to 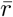. For the coat-independent case, we can calculate such a threshold by solving the equality at Eq. (8) for 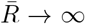 and different values of 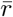 (see Fig. 4a). In particular, we can write *p*^⋆^ explicitly for 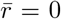 (see Appendix G) and, with the parameters reported in Table 1, we find 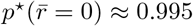 for 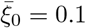, and 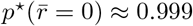 for 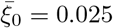.

Analogously, for a given inhibitor density *p* the inhibitor size 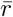 controls the range of particles for which the wrapping can initiate, and above a critical size 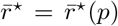 the process is completely blocked. Fig. 5 shows that for small protrusion sizes 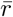, the selected size range is almost independent of moderate densities *p*, while this becomes relevant for larger protrusions.

**Figure 5:**
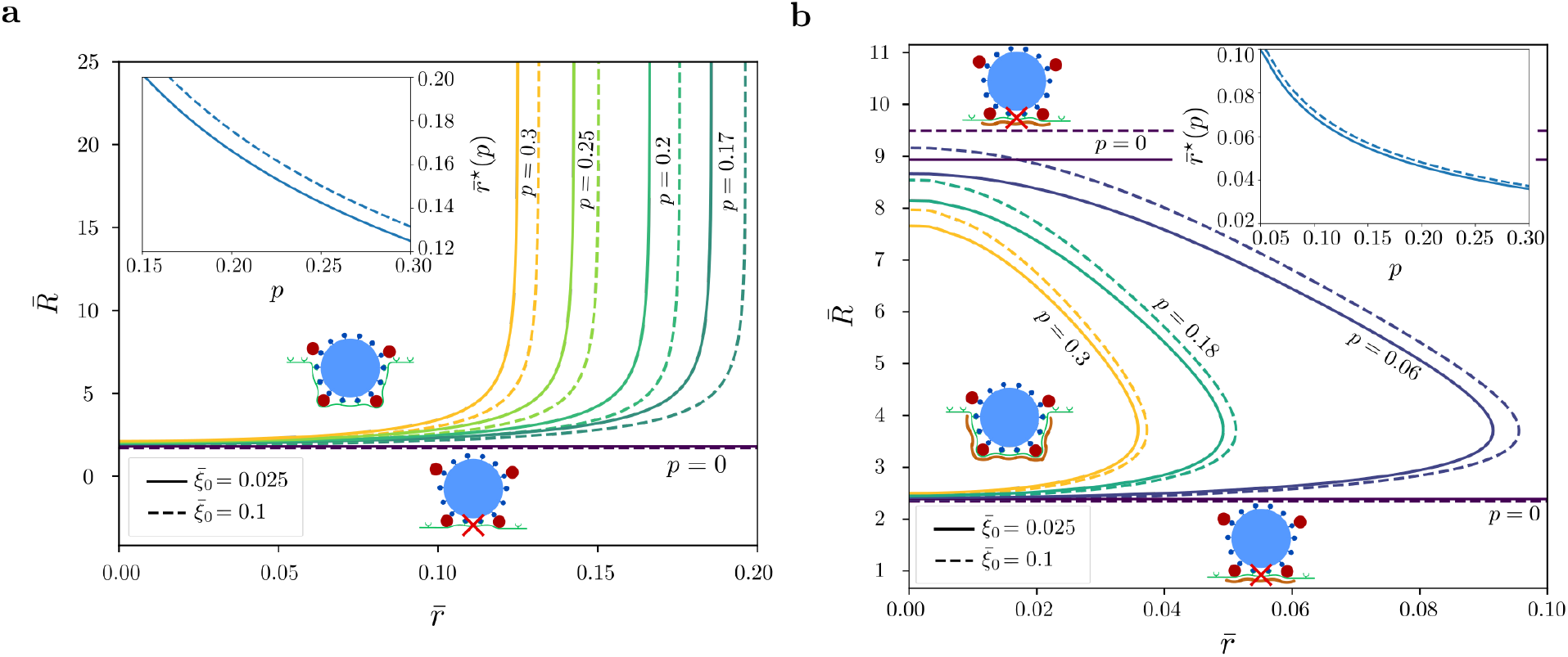
Regions of favorable endocytosis in the parameter space 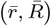 where 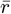 is the dimensionless obstacle sizes, 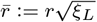 and 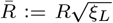 is the dimensionless particle radius. Region boundaries are determined from Eq. (8) for different fractions of inhibited ligands *p*, and ratios between ligand and receptor densities, 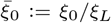. Insets show the critical inhibitor size *r*^⋆^ (*p*) above which endocytosis is completely blocked, as function of *p*. **(a)** Receptor-mediated endocytosis without protein coat for model parameters *d* = 1, *ē*_RL_ = 15, 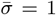, *ξ*_*L*_ = 5 ·10^−3^ nm^−2^, and 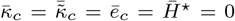. **(b)** Clathrin-mediated endocytosis for model parameters *d* = 1, *ē*_RL_ = 15, 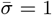, *ξ*_*L*_ = 5 ·10^−3^ nm^−2^, 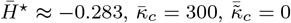, and *ē*_*c*_ = 23.

In both Fig. 4b and Fig. 5b, the range of radii [*R*_min_, *R*_max_] is asymmetric with respect to the preferred radius of curvature of the coat *R*^⋆^. This is due to the cost of bending the cell membrane, which diverges for 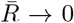 but vanishes for 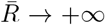, thus determining a lower bound but not an upper bound. It is the protein coat that introduces an upper bound in the particle size, because of its spontaneous curvature.

Finally, similar results follow by varying the indentation parameter *d* of the spherical inhibitors, while keeping fixed their size 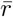. This corresponds to spherical inhibitors that protrude only partially, for *d* < 1, or that are raised above the particle surface, for *d >* 1 (see Fig. 3). Indeed, the elastic energy increases both by enlarging and by lifting the inhibitors, due to bending and tension. Thus, different geometries are equivalent in terms of the necessary condition for the wrapping given by Eq. (8). Fig. 6 shows contour plots of 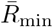 in the space of parameters that define the geometry of the protruding inhibitor, 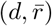, for the case of coat-independent endocytosis and for a fixed inhibitor density. As expected, the minimum radius is maximum when both the indentation and the size of the inhibitor are maximum, since such a configuration raises the cost of bending the cell membrane.

**Figure 6:**
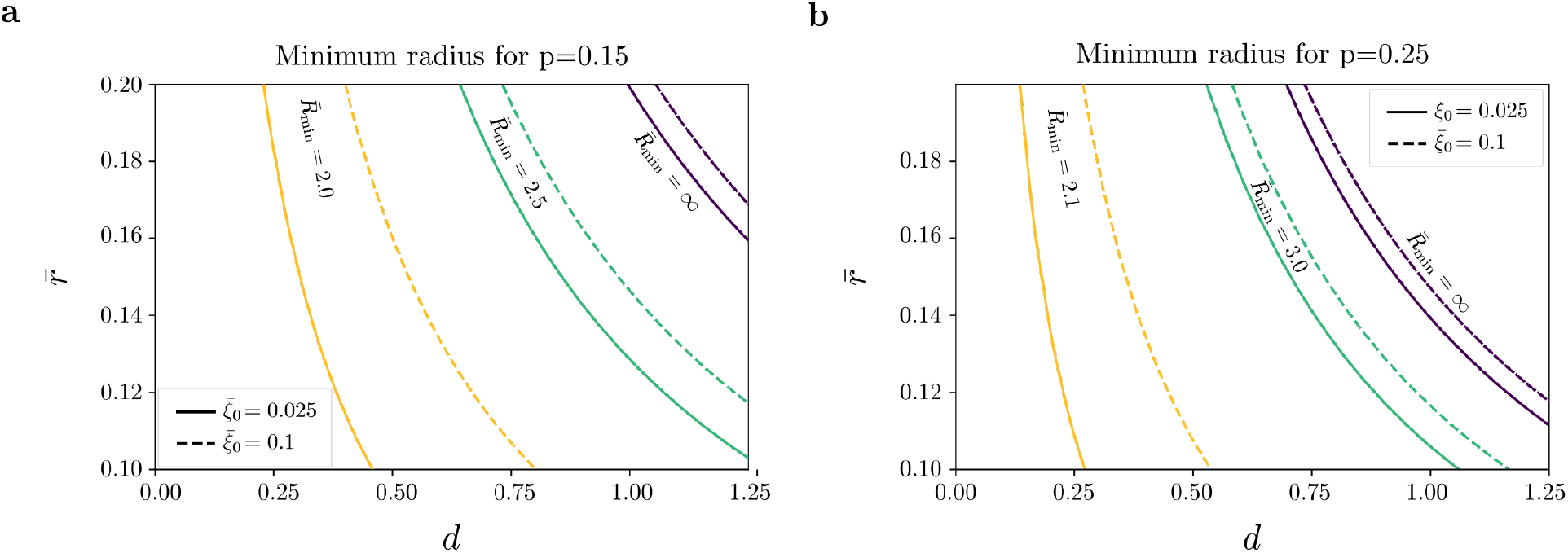
Level curves of the dimensionless minimum particle radius 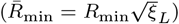 for favorable endocytosis as a function of the dimensionless radius of the inhibitor, 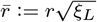, and *d*. The indentation depth of the protrusion in the membrane, from the surface of the particle, is *δ* = 2*dr* (see Fig. 2). Endocytosis is completely blocked by protrusions with parameters in the region on the right of the curves for 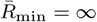. The results are for different values of the receptor density 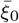 and the fraction of inhibited ligands *p*. Model parameters are *ē*_RL_ = 15, 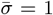, *ξ*_*L*_ = 5 ·10^−3^ nm^−2^, 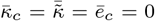 (no protein coat), and **(a)** *p* = 0.15 **(b)** *p* = 0.25.

## Discussion and conclusions

We found that the cost of bending the cell membrane and the protein coat, if present, is a determining factor for the inhibition of particle wrapping during the process of endocytosis. For the case of clathrin-mediated endocytosis, the cost of bending the cell membrane implies the existence of a minimum radius, while a maximum radius arises from the cost of bending the clathrin coat, due to its preferred curvature.

In the presence of inhibitors, the wrapping process depends on the radius and indentation depth of their protrusion, as well as their density. This reduces the size range of particles for which the wrapping process can initiate. Moreover, for a given inhibitor density, the uptake is completely blocked when the radius and indentation depth of the inhibitor are above given thresholds (see Fig. 6).

We analyzed the case of a spherical particle with uniformly distributed protruding inhibitors. Real biological systems are expected to exhibit random distribution of inhibitors/protrusions, hence our model provides a coarse-grained analysis over a large population of particles/viruses.The current study provides a simple way to estimate the efficacy of ligand-inhibitors in selectively blocking endocytosis in the presence and in the absence of protein (clathrin) coats. Quantitative validation of our findings is difficult, due to the lack of experiments on the morphology considered in this study. However, when specialized to the case of spherical particles without protrusions, our results agree with experimental observations on clathrin-mediated endocytosis (Rejman et al., 2004), and are consistent with previous findings in the absence of clathrin (Gao et al., 2005).

Given our focus on the (un)favorable conditions for the initiation of the wrapping process, we neglected elastic deformations outside the wrapping zone. We also neglected the energetic contribution of cortical actin, which might help the formation of clathrin-coated vesicles (Boulant et al., 2011; Collins et al., 2011). However, future research can investigate the role of all these processes by considering approaches available in the literature (Zhang et al., 2015; Yi and Gao, 2017; Hassinger et al., 2017).

In conclusion, it is worth emphasizing the potential implications of the present study. Our model provides a tool for the design of novel molecular strategies for the morphological control of endocytosis, with applications ranging from viral antibodies to engineered nanoparticles for targeted diagnostics and therapeutics. Indeed, our results suggest at least two strategies for targeted uptake. One possibility is to exploit overexpressed receptors on targeted cells to deliver nanoparticles that would not enter for lower receptor densities (this corresponds to nanoparticles between continuous and dashed lines in Fig. 4 and Fig. 5). Another possibility is to design a particle with protrusions that can bind only to targeted cells, such that more restrictive conditions would apply to enter other cells.

## Contributions

M.B. designed the research; D.A. performed the research; D.A., G.J.E, and M.B. wrote the paper.

## Acknowledgments

The authors gratefully acknowledge the useful discussions with Dr. Don Sin (Saint Paul Hospital) and Masahiro Niikura (Simon Fraser University). M.B. and G.J.E. acknowledge the financial support from the New Frontiers in Research Funds - Exploration (NFRFE-2018-00730); M.B. acknowledges the financial support from the Natural Sciences and Engineering Research Council of Canada (NSERC) (RGPIN-2017–04464, and ALLRP554607-20); D.A. acknowledges the financial support from the Michael Smith Foundation for Health Research (RT-2021-1954).

## Appendices

### Appendix A. Spacing between particle ligands

For a large number *N* ≈ 4*πR*^2^*ξ*_*L*_ of ligands, we approximate the sphere of radius *R* with a polyhedron with *H* ≈*N*/3 hexagons of area 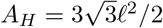. By comparing the surface areas of the polyhedron and the sphere, namely,

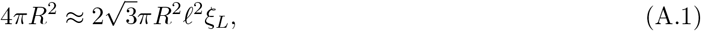

we arrive at 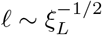, which we adopt in our model as the spacing between two ligands.

### Appendix B. The maximum number of inhibitors

In this study we assume that the wrapping process of the particle with inhibitors is driven by receptorligand binding around the protrusion. This condition is possible for the fraction of inhibited ligands *p* that is below a critical value *p*_max_, to assure the presence of a sufficient number of active ligands. Each protrusion is associated with a surface area of ∼ 2*πR*^2^ (1 −cos *ℓ/R*) on the sphere. In the limit case of *p* = *p*_max_, there are at most

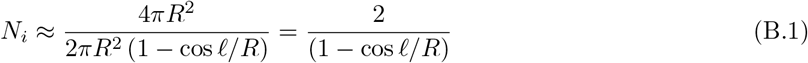

inhibitors for *N* ≈ 4*πR*^2^*ξ*_*L*_ ligands. Because *p* = *N*_*i*_*/N*, from Eq. (B.1), we define

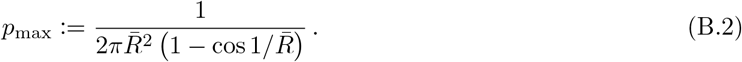

In the limit of a large dimensionless radius 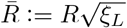, we get *p*_max_ ≈ 1*/π*. We observe that this is approximately in agreement with the hexagonal distribution, for which each ligand is adjacent to 3 protrusions, such that each protrusion is associated with 2 ligands and *p*_max_ ≈ 1/3.

### Appendix C. Condition for endocytosis

In this section we derive the condition for endocytosis of a particle with protruding inhibitors. The free energy of the cell membrane is the sum of the surface free energy density integrated over the contact region 𝒮_*w*_, and over the remaining area of the membrane, 𝒮 \ 𝒮_*w*_, giving

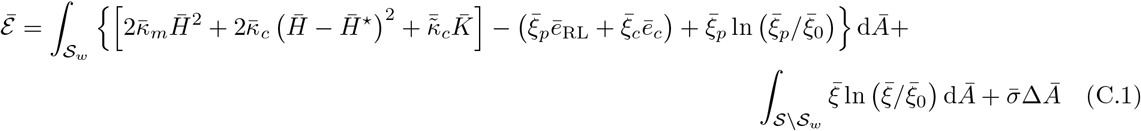

with the notation introduced in the main text. In particular, we recall that *ξ*_*p*_ is the receptor density on the cell membrane in the contact region given by Eq. (1).

Splitting the wrapping region 𝒮_*w*_ into representative regions, 𝒮_*p*_, wrapping one single protrusion, and the remaining surface adhering to the spherical part of the particle, from Eq. (1) and Eq. (C.1) we have

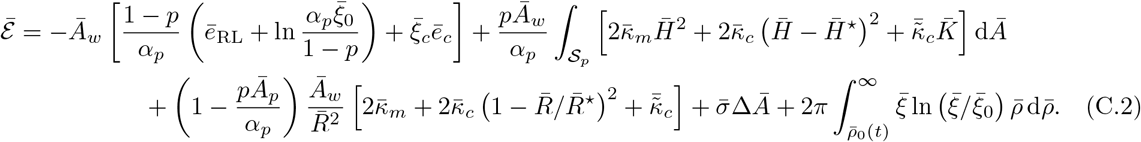

Here we used the polar coordinate *ρ* (given the axial symmetry of the problem) to rewrite the integral over the unwrapped (flat) region 𝒮 \ 𝒮_*w*_ with *ρ*_0_ the radius of the projection of 𝒮_*w*_ on the plane, 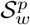(see Fig. 2a).

During wrapping the contact area *A*_*w*_ increases with time and *ρ*_0_ varies accordingly. For protrusions of small size, *r*, with respect to the particle radius, *R*, we have

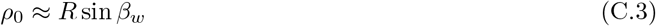

where *β*_*w*_ is the contact (wrapping) angle (Fig. 2a). Moreover,

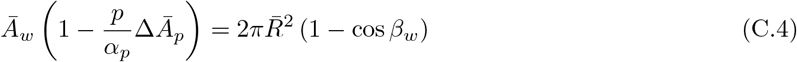

where 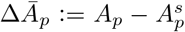 is the difference between the surface area of 𝒮_*p*_ (*A*_*p*_) and the one of its projection on the spherical particle 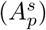. Combining Eq. (C.3) and Eq. (C.4), we arrive at

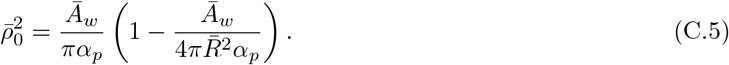

Following Freund and Lin (2004), we assume that receptors diffuse through the cell membrane according to the second Fick’s law giving the rate of change of receptor density as 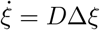, where *D* is the diffusion coefficient and a dot denotes the time derivative. In axial symmetry this equation reads as

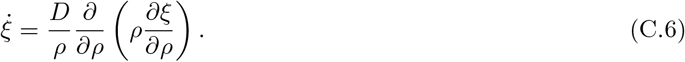

Rewriting then Eq. (C.6) in dimensionless form we have

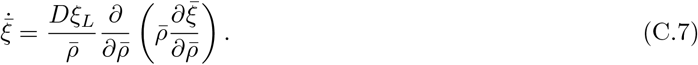

Moreover, by imposing the conservation of cell receptors, we get

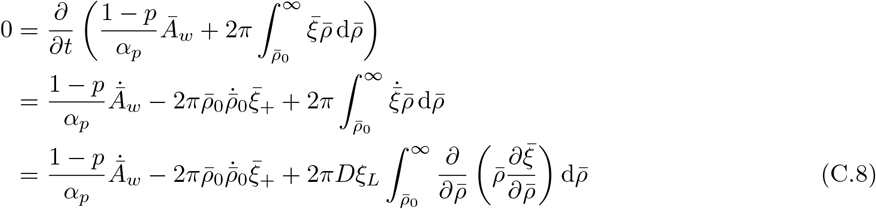

whence

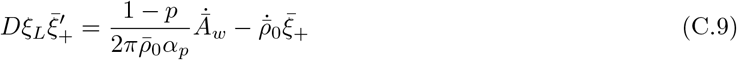

where 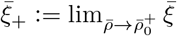 and 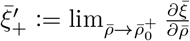 are the limit values of the cell receptor density and its radial derivative at the front of the wrapping region, respectively.

Finally, by taking the time derivative of the free energy (C.2), plugging in Eq. (C.7), integrating by parts, and exploiting Eq. (C.9) together with the time derivative of Eq. (C.5) and the approximation 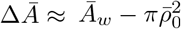, we arrive at

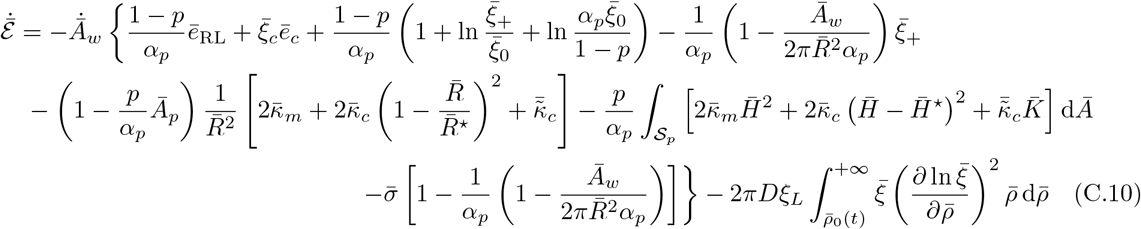

where the last integral term represents the rate of energy consumed during receptor transport (Freund and Lin, 2004). By assuming that the rate of the free energy reduction is greater than or equal to the one of energy dissipation due to receptor transport, we derive the inequality

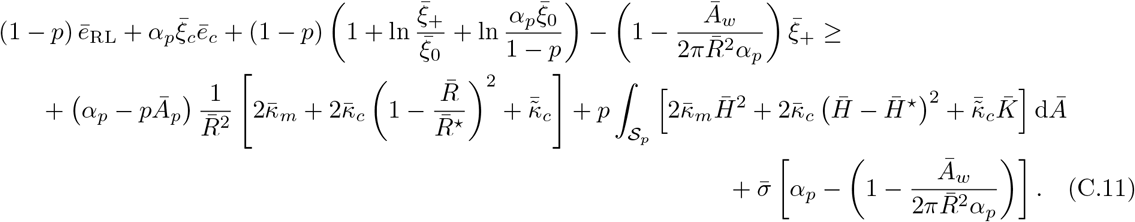

Finally, by approximating 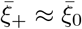 and 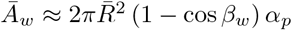, we get

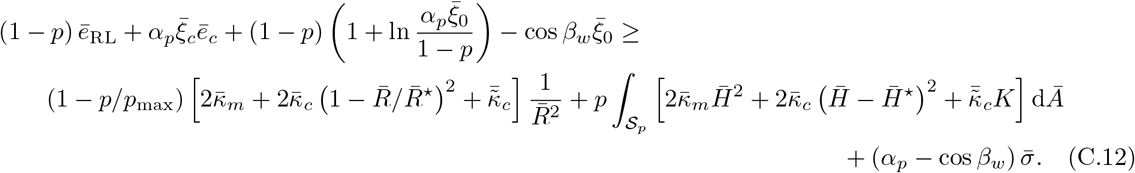

Eq. (8) follows by setting *β*_*w*_ = 0 in Eq. (C.12), and it reduces to Eq. (7) for *p* = 0.

### Appendix D. The critical spontaneous curvature

Eq. (7) provides a necessary condition for initiating wrapping of a spherical particle. This is a quadratic inequality in the dimensionless radius 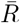, namely,

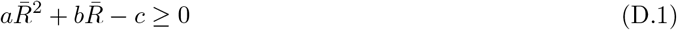

where 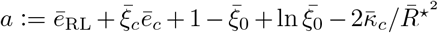, and 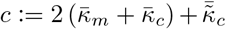. Thus, the presence of a protein coat with a spontaneous curvature can determine a maximum value 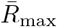 for the particle radius if *a* < 0, *i*.*e*. if the spontaneous radius of curvature satisfies

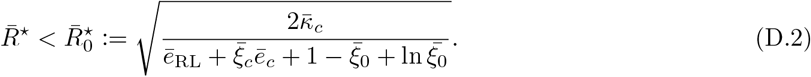

In passing, we notice that we implicitly assumed that 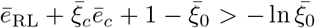, otherwise the binding energy would be insufficient to drive the wrapping.

### Appendix E. The obstacle problem

We determine the configuration of the membrane in a region associated with a protrusion, 𝒮_*p*_, as solution to the obstacle problem (*i*.*e*. the minimization of the free energy of the membrane within the limits of compatibility)

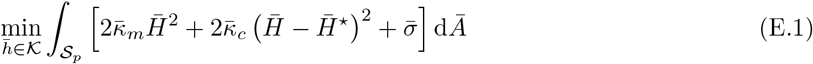

where 𝒦 is the set of all admissible configurations for the cell membrane wrapping the inhibitor and being clamped to the particle at the boundary ∂*𝒮*_*p*_. In Eq. (E.1) we dropped the saddle-splay term of the protein coat because the prescribed boundary conditions force the geodesic curvature at the boundary so that integral of the Gaussian curvature amounts to a constant, in view of the Gauss-Bonnet theorem.

We adopt the Monge parametrization to describe both the membrane and the obstacle and, by assuming axial symmetry of the problem, we introduce the corresponding dimensionless heights 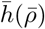 and 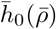 as functions of the dimensionless polar coordinate 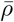. Thus, the problem reads as

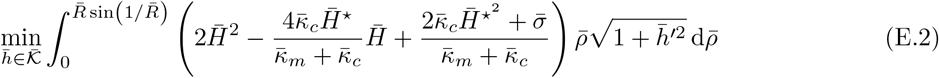

with the membrane configuration 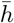 belonging to the set of sufficiently regular height functions that fulfill the compatibility constraints, *i*.*e*.,

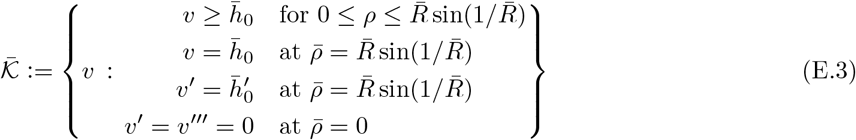

where primes denote differentiation with respect to 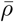. In particular, we define the obstacle as

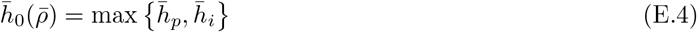

where 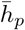 is the height function of the particle, and 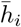 is the height function of the inhibitor, namely

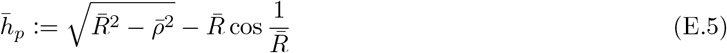

and

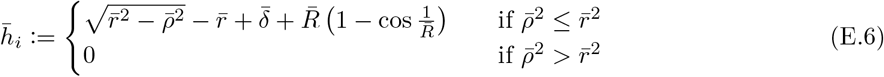

where 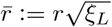 is the dimensionless radius of the spherical protrusion and 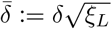 is the dimensionless indentation *δ* = 2*dr* (see Fig. 3).

In terms of the adopted parametrization, the mean curvature of the membrane having height 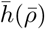 is given by

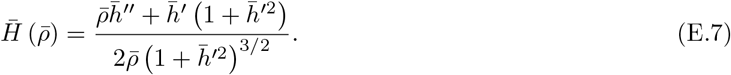

The obstacle problem (E.2)-(E.4) depends only on the three dimensionless parameters that define the specific geometry of the obstacle, *i*.*e*., 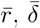, and 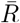, and the two dimensionless parameters 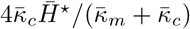 and 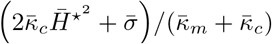. This problem is a variational inequality and we solve it numerically by using the penalty method (*e*.*g*., see Glowinski (1984) and references therein), namely, we minimize the penalized functional

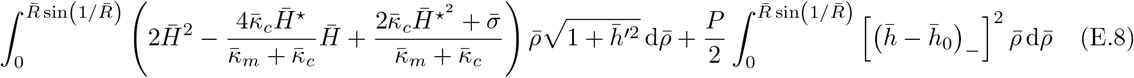

where (·)_−_:= min {0, ·} is the “negative part” function and *P* ≫ 1 is the penalty parameter.

We derive the weak form of the problem as a system of the two following variational equations, *i*.*e*.,

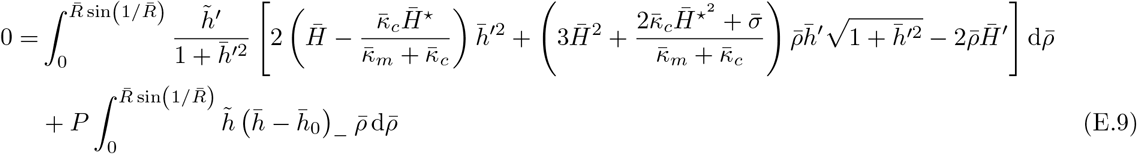

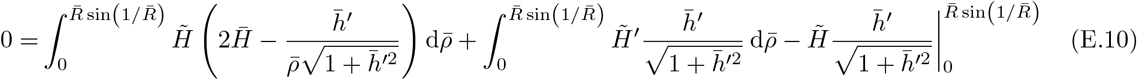

where 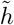 and 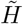 are the test functions.

We implemented and solved the weak formulation (E.9)-(E.10) via finite element analysis based on the FEniCS Project Version 2019.1.0 (Logg et al., 2012). For our numerical simulations we used a penalty parameter *P* = 10^8^. The Python code is available at https://github.com/daniele-agostinelli/ObstacleProblem.git.

### Appendix F. Single-spaced and double-spaced configurations

Throughout this study we assumed that the cellular membrane wrapping around an inhibitor binds to the closest active ligands located at distance *ℓ* from the inhibitor (single-spaced configuration). However, other configurations might be more energetically convenient, for example one in which the membrane binds to active ligands located at a distance 2*ℓ* (double-spaced configuration). If each inhibitor is associated with an area corresponding to *n* times the angle *ℓ/R*, there are at most 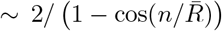 inhibitors. By comparing this number with 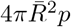, we get the relationship

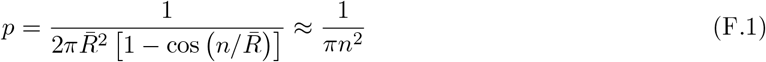

where the last approximation holds for 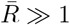 (large particle or high ligand density).

From the estimate (F.1), a configuration with a doubled angle (double-spaced) is possible for *p* < 0.08. In this case we can compare the energies of the single-spaced (*n* = 1) and double-spaced (*n* = 2) configurations shown in Fig. F.7. The latter is convenient if

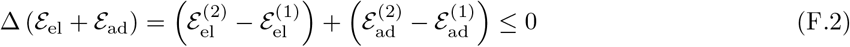

with ^(1)^ indicating single-spaced configuration and ^(2)^ double-spaced.

**Figure F.7:**
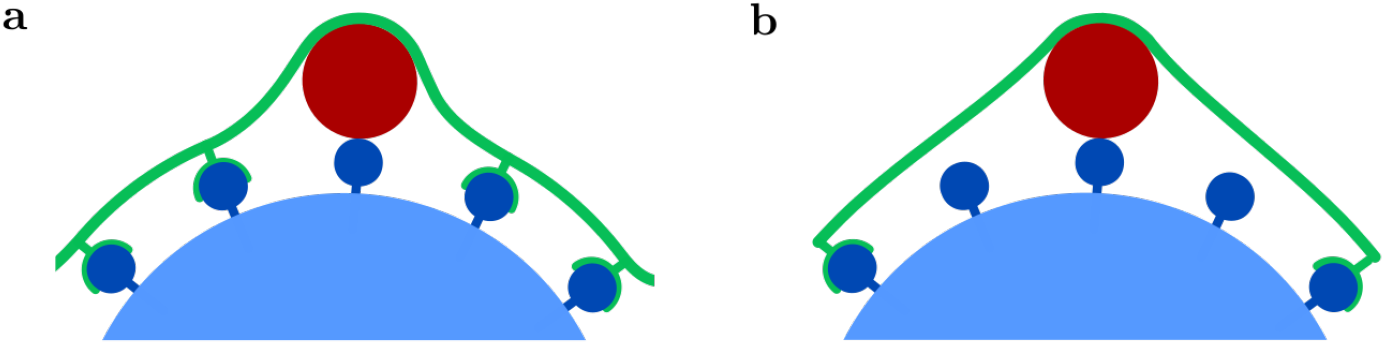
Two-dimensional illustration of the **(a)** single-space (*n* = 1) and **(b)** double-spaced (*n* = 2) configurations. For a low fraction of inhibited ligands, *p* ∼ 0.1, the former might be energetically more expensive than the latter, where cell receptors do not bind to the ligands that are proximal to the protrusion.

Fig. F.8 shows the domains of the two configurations in the parameter space 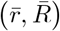, for selected values of the model parameters. Fig. F.8a reports the case of receptor-mediated endocytosis, in the absemce of protein coats, while Fig. F.8b reports the case of protein coat. In the latter case, compared to the former, we observe higher sensitivity to 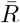, while receptor density 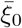 is less influential.

**Figure F.8:**
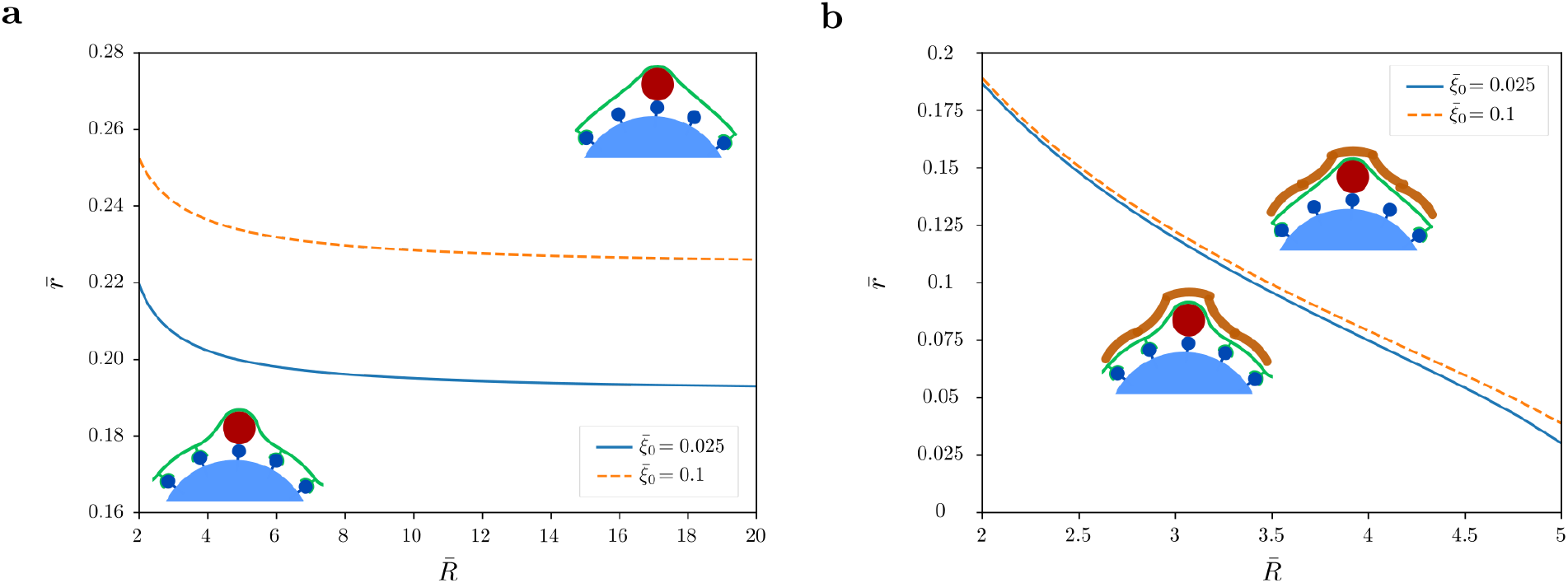
Comparison between the energetic cost of single-spaced and double-spaced configurations: larger protrusions (increasing 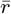) leads to higher bending cost, which might (not) be compensated by the enthalpic gain due to receptor-ligand bindings, thus resulting in a critical threshold of the inhibitor size, 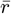. The parameter space 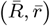 is split into regions where one configuration is more convenient than the other one, as indicate in the plots (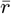 above the line provides double-spaced). We report the boundaries of these regions for different values of the ratio between receptors and ligands, 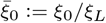, for the case of **(a)** receptor-mediated endocytosis without any protein coat 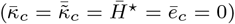 and **(b)** clathrin-mediated endocytosis for 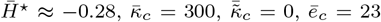. We set *d* = 1 and other model parameters as *ē*_RL_ = 15, 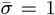, *ξ*_*L*_ = 5 ·10^−3^ nm^−2^, *p* ≈ 0.08.

### Appendix G. The effect of inhibiting ligands without protrusions

In this appendix we examine the effect of inhibiting particle ligands without protrusions (*p >* 0 and 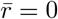). In this case Eq. (8) yields

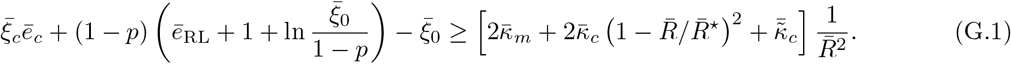

By assuming that 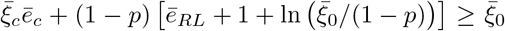, the presence of clathrin provides both a minimum and a maximum value for the particle radius if the spontaneous radius of curvature satisfies

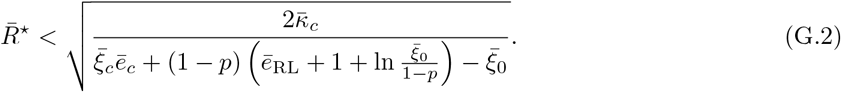

If clathrin is absent or the spontaneous radius of curvature is above such a threshold, there exists only a minimum particle radius. In particular, in the absence of clathrin, we get

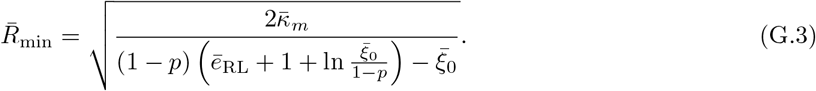

Since 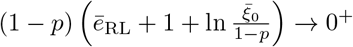 for *p* → 1^−^, the denominator vanishes at *p* = *p*^⋆^ where *p*^⋆^ < 1. More precisely, for the relevant parameter values,

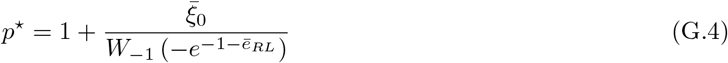

where *W*_*k*_(·) denotes the *k*^th^ branch of the Lambert W-Function. Then 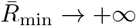 for *p* → *p*^⋆−^. We observe that the critical fraction 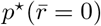 is above 97% for the parameters used in this study, and 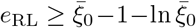, which is the minimum receptor-ligand binding energy for having sufficient driving force. This implies that, in the absence of clathrin, inhibiting ligands without additional bending penalty is an inefficient method of blocking the wrapping, since almost all ligands should be inhibited.

## Notes

### Competing Interest Statement

The authors have declared no competing interest.

## References

Banerjee, A., Berezhkovskii, A., Nossal, R., 2012. Stochastic model of clathrin-coated pit assembly. Biophysical Journal 102, 2725–2730. URL: https://www.sciencedirect.com/science/article/pii/S0006349512005632, doi:https://doi.org/10.1016/j.bpj.2012.05.010.

Boulant, S., Kural, C., Zeeh, J.C., Ubelmann, F., Kirchhausen, T., 2011. Actin dynamics counteract membrane tension during clathrin-mediated endocytosis. Nature cell biology 13, 1124–1131. URL: https://doi.org/10.1038/ncb2307, doi:10.1038/ncb2307.

Canham, P., 1970. The minimum energy of bending as a possible explanation of the biconcave shape of the human red blood cell. Journal of Theoretical Biology 26, 61–81. URL: https://www.sciencedirect.com/science/article/pii/S0022519370800327, doi:https://doi.org/10.1016/S0022-5193(70)80032-7.

Collins, A., Warrington, A., Taylor, K.A., Svitkina, T., 2011. Structural organization of the actin cytoskeleton at sites of clathrin-mediated endocytosis. Current Biology 21, 1167–1175. URL: https://www.sciencedirect.com/science/article/pii/S0960982211006063, doi:10.1016/j.cub.2011.05.048.

Derényi, I., Jülicher, F., Prost, J., 2002. Formation and interaction of membrane tubes. Phys. Rev. Lett. 88, 238101. URL: https://link.aps.org/doi/10.1103/PhysRevLett.88.238101, doi:10.1103/PhysRevLett.88.238101.

Deserno, M., 2004. Elastic deformation of a fluid membrane upon colloid binding. Phys. Rev. E 69, 031903. URL: https://link.aps.org/doi/10.1103/PhysRevE.69.031903, doi:10.1103/PhysRevE.69.031903.

Flint, J., Racaniello, V., Rall, G., Skalka, A., Enquist, L., 2015. Principles of virology, Volume 1: Molecular biology. 4th ed., ASM Press.

Freund, L., Lin, Y., 2004. The role of binder mobility in spontaneous adhesive contact and implications for cell adhesion. Journal of the Mechanics and Physics of Solids 52, 2455–2472. URL: https://www.sciencedirect.com/science/article/pii/S0022509604001012, doi:https://doi.org/10.1016/j.jmps.2004.05.004.

Frey, F., 2019. Physical models for uptake processes at the cell membrane. Ph.D. thesis.

Gao, H., Shi, W., Freund, L., 2005. Mechanics of receptor-mediated endocytosis. Proceedings of the National Academy of Sciences 102, 9469–9474. URL: https://www.pnas.org/content/102/27/9469, doi:10.1073/pnas.0503879102, arXiv:https://www.pnas.org/content/102/27/9469.full.pdf.

Glowinski, R., 1984. Generalities on Elliptic Variational Inequalities and on Their Approximation. Springer Berlin Heidelberg, Berlin, Heidelberg. pp. 1–26. URL: https://doi.org/10.1007/978-3-662-12613-4_1, doi:10.1007/978-3-662-12613-4_1.

Hassinger, J., Oster, G., Drubin, D., Rangamani, P., 2017. Design principles for robust vesiculation in clathrin-mediated endocytosis. Proceedings of the National Academy of Sciences 114, E1118–E1127. URL: https://www.pnas.org/content/114/7/E1118, doi:10.1073/pnas.1617705114, arXiv:https://www.pnas.org/content/114/7/E1118.full.pdf.

Helfrich, W., 1973. Elastic properties of lipid bilayers: Theory and possible experiments. Zeitschrift für Naturforschung C 28, 693–703. URL: https://doi.org/10.1515/znc-1973-11-1209, doi:10.1515/znc-1973-11-1209.

Heuser, J., 1980. Three-dimensional visualization of coated vesicle formation in fibroblasts. Journal of Cell Biology 84, 560–583. URL: https://doi.org/10.1083/jcb.84.3.560, doi:10.1083/jcb.84.3.560, arXiv:https://rupress.org/jcb/article-pdf/84/3/560/1401003/560.pdf.

Kumar, G., Sain, A., 2016. Shape transitions during clathrin-induced endocytosis. Phys. Rev. E 94, 062404. URL: https://link.aps.org/doi/10.1103/PhysRevE.94.062404, doi:10.1103/PhysRevE.94.062404.

Lauster, D., Klenk, S., Ludwig, K., Nojoumi, S., Behren, S., Adam, L., Stadtmüller, M., Saenger, S., Zimmler, S., Hönzke, K., et al., 2020. Phage capsid nanoparticles with defined ligand arrangement block influenza virus entry. Nature nanotechnology 15, 373–379. URL: https://doi.org/10.1038/s41565-020-0660-2, doi:10.1038/s41565-020-0660-2.

Lewis, B., Engelman, D., 1983. Lipid bilayer thickness varies linearly with acyl chain length in fluid phosphatidylcholine vesicles. Journal of Molecular Biology 166, 211–217. URL: https://www.sciencedirect.com/science/article/pii/S0022283683800072, doi:10.1016/S0022-2836(83)80007-2.

Logg, A., Mardal, K.A., Wells, G., 2012. Automated solution of differential equations by the finite element method: The FEniCS book. volume 84. Springer Berlin Heidelberg, Berlin, Heidelberg. URL: https://doi.org/10.1007/978-3-642-23099-8, doi:10.1007/978-3-642-23099-8.

Marsh, M., Helenius, A., 2006. Virus entry: Open sesame. Cell 124, 729–740. URL: https://www.sciencedirect.com/science/article/pii/S0092867406001826, doi:https://doi.org/10.1016/j.cell.2006.02.007.

Mashl, R., Bruinsma, R., 1998. Spontaneous-curvature theory of clathrin-coated membranes. Biophysical Journal 74, 2862–2875. URL: https://www.sciencedirect.com/science/article/pii/S0006349598779937, doi:https://doi.org/10.1016/S0006-3495(98)77993-7.

Morris, C., Homann, U., 2001. Cell surface area regulation and membrane tension. The Journal of membrane biology 179, 79–102. URL: https://doi.org/10.1007/s002320010040, doi:10.1007/s002320010040.

Nossal, R., 2001. Energetics of clathrin basket assembly. Traffic 2, 138–147. URL: https://onlinelibrary.wiley.com/doi/abs/10.1034/j.1600-0854.2001.020208.x, doi:https://doi.org/10.1034/j.1600-0854.2001.020208.x, arXiv:https://onlinelibrary.wiley.com/doi/pdf/10.1034/j.1600-0854.2001.020208.x.

Pedersen, J., 1993. Structure of clathrin-coated vesicles from small-angle scattering experiments. European biophysics journal 22, 79–95. URL: https://doi.org/10.1007/BF00196913, doi:10.1007/BF00196913.

Rejman, J., Oberle, V., Zuhorn, I., Hoekstra, D., 2004. Size-dependent internalization of particles via the pathways of clathrin- and caveolae-mediated endocytosis. Biochemical Journal 377, 159–169. URL: https://doi.org/10.1042/bj20031253, doi:10.1042/bj20031253, arXiv:https://portlandpress.com/biochemj/article-pdf/377/1/159/712500/bj3770159.pdf.

Saleem, M., Morlot, S., Hohendahl, A., Manzi, J., Lenz, M., Roux, A., 2015. A balance between membrane elasticity and polymerization energy sets the shape of spherical clathrin coats. Nature communications 6, 1–10. URL: https://doi.org/10.1038/ncomms7249, doi:10.1038/ncomms7249.

Steigmann, D., 2017. The role of mechanics in the study of lipid bilayers. volume 577 of CISM International Centre for Mechanical Sciences. Springer, Cham. URL: https://link.springer.com/book/10.1007%2F978-3-319-56348-0, doi:https://doi.org/10.1007/978-3-319-56348-0.

Stoeber, M., Schellenberger, P., Siebert, C., Leyrat, C., Helenius, A., Grünewald, K., 2016. Model for the architecture of caveolae based on a flexible, net-like assembly of cavin1 and caveolin discs. Proceedings of the National Academy of Sciences 113, E8069–E8078. URL: https://www.pnas.org/content/113/50/E8069, doi:10.1073/pnas.1616838113, arXiv:https://www.pnas.org/content/113/50/E8069.full.pdf.

Traub, L., 2011. Regarding the amazing choreography of clathrin coats. PLOS Biology 9, 1–5. URL: https://doi.org/10.1371/journal.pbio.1001037, doi:10.1371/journal.pbio.1001037.

Vigers, G., Crowther, R., Pearse, B., 1986. Location of the 100 kd-50 kd accessory proteins in clathrin coats. The EMBO Journal 5, 2079–2085. URL: https://www.embopress.org/doi/abs/10.1002/j.1460-2075.1986.tb04469.x, doi:https://doi.org/10.1002/j.1460-2075.1986.tb04469.x, arXiv:https://www.embopress.org/doi/pdf/10.1002/j.1460-2075.1986.tb04469.x.

Yi, X., Gao, H., 2017. Kinetics of receptor-mediated endocytosis of elastic nanoparticles. Nanoscale 9, 454—463. URL: https://doi.org/10.1039/c6nr07179a, doi:10.1039/c6nr07179a.

Yi, X., Shi, X., Gao, H., 2014. A universal law for cell uptake of one-dimensional nanomaterials. Nano Letters 14, 1049–1055. URL: https://doi.org/10.1021/nl404727m, doi:10.1021/nl404727m, arXiv:https://doi.org/10.1021/nl404727m. pMID: 24459994.

Zhang, S., Gao, H., Bao, G., 2015. Physical principles of nanoparticle cellular endocytosis. ACS Nano 9, 8655–8671. URL: https://doi.org/10.1021/acsnano.5b03184, doi:10.1021/acsnano.5b03184, arXiv:https://doi.org/10.1021/acsnano.5b03184. pMID: 26256227.

